# Synthetic dysmobility screening reveals a phosphorylation-independent facet of the SLK-Ezrin-F-actin signaling cascade

**DOI:** 10.1101/2024.07.23.604488

**Authors:** Hsuan-Chao Lin, Yun-Yu Lin, Ling-Ying Wei, Jia-Ming Zhang, Pei-Ju Tsai, Yu-Chiao Lin, Jean-San Chia, Feng-Chiao Tsai

## Abstract

This study investigates the role of Ste20-like kinase (SLK) in cytoskeletal dynamics, focusing on its interaction with Ezrin and actin remodeling, independent of phosphorylation processes. We utilized HaCaT to explore the effects of SLK and Ezrin knockdown, as well as the application of specific inhibitors on the organization of actin structures. Our findings reveal that both SLK and Ezrin significantly influence the architecture of actin cytoskeletons with differential impacts observed between protein knockdown and dephosphorylation events. Additionally, our results suggest novel, phosphorylation-independent pathways through which Ezrin modulates actin dynamics, potentially indicating alternative, non-enzymatic roles for Ezrin in cytoskeletal integrity.

## Introduction

The Ste20-like kinase (SLK) emerges as a multifaceted regulatory protein with significant implications in cellular dynamics and signaling pathways. Primarily associated with the microtubule network, SLK plays a crucial role in modulating cytoskeletal architecture, particularly in the context of cell adhesion and migration (*1-5*). Our screening data also reveal that SLK significantly enhances both the speed and coordination of cell migration (*6*). Prior studies have demonstrated that SLK directly phosphorylates and activates ERM (ezrin/radixin/moesin) proteins, facilitating their localization at the cell cortex. Specifically, SLK targets ezrin at threonine 567 (T567), a critical regulatory site (*7*), which promotes ezrin’s activation and subsequent association with the plasma membrane (*8, 9*). Active RhoA directly binds to the C-terminal coiled-coil domain of SLK, leading to SLK activation (*10*). This interaction promotes SLK-mediated phosphorylation of ERM proteins, which are crucial regulators of the actin cytoskeleton and cell cortex. The RhoA-SLK-ERM signaling axis represents a previously unrecognized mechanism by which RhoA influences cellular architecture and shape (*11*).

Our study delineates distinct pathways through which SLK and Ezrin influence the architecture of actin fibers, particularly emphasizing the regulatory cascades independent of Ezrin’s phosphorylation state. We observed that suppressing SLK or Ezrin not only disrupts normal actin structures but also does so in ways that are not solely a consequence of inhibited Ezrin phosphorylation. This suggests that both proteins orchestrate actin architecture through diverse, yet overlapping, pathways. Furthermore, we discovered that Ezrin modulates actin fibers in a phosphorylation-independent manner; its ability to reduce contractile actin fibers does not rely on its phosphorylation state, indicating that Ezrin can influence actin dynamics through alternative, non-enzymatic pathways. These findings highlight that the interaction between SLK and Ezrin in regulating actin structures extends beyond enzymatic actions, involving more complex, possibly structural or scaffolding functions that help maintain cytoskeletal integrity.

## Results

### Distinct impact of SLK and Ezrin knockdown on actin structures compared to Ezrin dephosphorylation

Our study began by investigating whether Ste20-like kinase (SLK) affects cellular behaviors, such as cell migration, through known mechanisms, particularly the phosphorylation of Ezrin and its impact on F-actin polymerization. Data from the Human Protein Atlas indicate that SLK expression is predominantly seen in epithelial cells, exhibiting higher expression levels in HaCaT cells (a keratinocyte line) compared to HUVEC cells (Human Umbilical Vein Endothelial Cells). Given the relative ease of culturing and genetic manipulation in HaCaT cells, coupled with the more pronounced morphological changes observed upon SLK knockdown in these cells, our experimental efforts have been concentrated on this cell line to further elucidate the role of SLK in cytoskeletal dynamics. Initial experiments using HaCaT cells with phalloidin staining revealed that knockdown of either SLK or Ezrin increases F-actin cortex meshwork expression. Notably, the effect of Ezrin knockdown was distinct, enhancing the membrane-tethered actin to a lesser extent while preserving contractile stress fibers (Fig. 1, A and B). To confirm the shRNA knockdown results, we used specific inhibitors: Cpd31 for SLK and NSC305787 or NSC668394 for Ezrin (*12, 13*). Treatment with these inhibitors similarly increased the membrane-tethered actin, yet resulted in morphologies distinct from those seen with Ezrin knockdown alone (Fig. 1, C and D). HUVEC cells exhibited similar outcomes; the knockdown of either SLK or Ezrin resulted in an increased presence of membrane-tethered actin and contractile stress fibers (Fig. 1, E and F). According to the Brownian ratchet hypothesis, random molecular motion can be harnessed to generate directed mechanical work, as molecular fluctuations are rectified into unidirectional movement by structural asymmetries in molecular machinery (*14-17*). We are investigating whether cell wrinkling can be attributed to the Brownian ratchet hypothesis. Additionally, we aim to determine whether Ezrin inhibitors contribute to this phenomenon by hypothesizing that increased robustness in the cell cortex could induce slight membrane curvature. Time-dependent treatment with these inhibitors at low concentrations for 25 hours consistently resulted in low levels of phosphorylated ERM (p-ERM), with the bent cell membrane phenotype remaining stable over time (fig. S1). Our findings reveal that both SLK and Ezrin significantly influence the architecture of actin cytoskeletons, with differential impacts observed between protein knockdown and dephosphorylation events.

**Fig 1.**
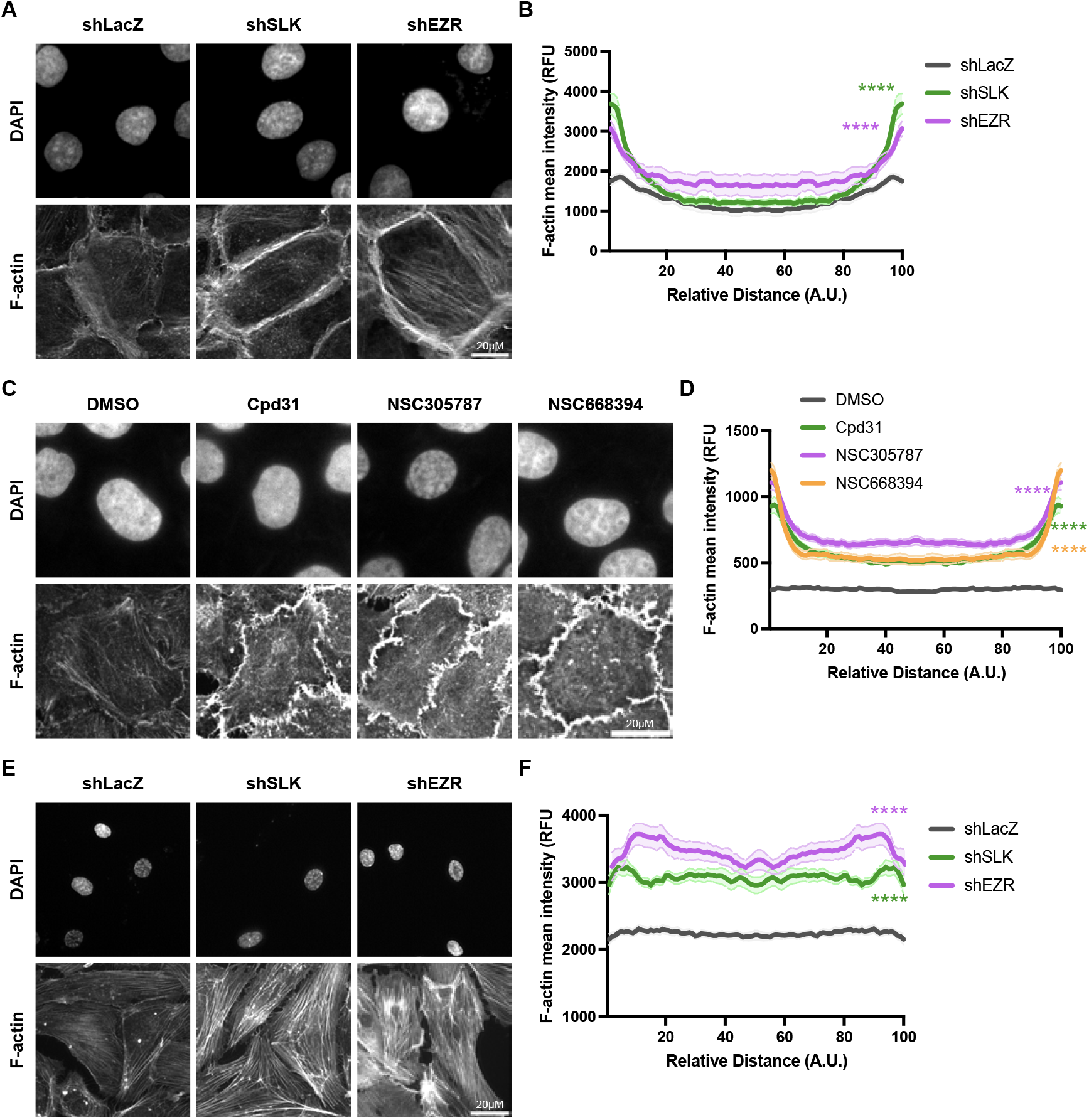
SLK or Ezrin knockdown altered actin structures distinctly from there by the suppression of Ezrin phosphorylation. (A)(B) Knockdown of SLK leads to an elevation in contractile actin fibers, while knockdown of EZR results in an increase in stress fibers. (C)(D) The suppression of Ezrin phosphorylation leads to a reduction in contractile actin fibers and an elevation in membrane-tethered actin. (E)(F) HUVEC exhibited similar outcomes to HaCaT cells, albeit with a greater abundance of stress fibers. Actin signal results were calculated as the average of 12 different lines drawn through each cell at various angles. One-way analysis of variance (ANOVA) and Scheffe’s/Dunn post hoc comparison. Mean ± s.e.m.

### Double inhibition experiments revealed a novel phosphorylation-independent SLK-EZR pathway on actin remodeling

Based on our observations, we hypothesized that non-phosphorylated Ezrin could influence the distribution or formation of F-actin. To explore this hypothesis, we conducted experiments involving the knockdown of Ste20-like kinase (SLK) alone and in conjunction with Ezrin inhibitors. We observed that the simultaneous knockdown of SLK and Ezrin exhibited phenotypes typical of each individual knockdown, implying an influence of non-phosphorylated Ezrin on F-actin structure (Fig. 2, A and B).

**Fig 2.**
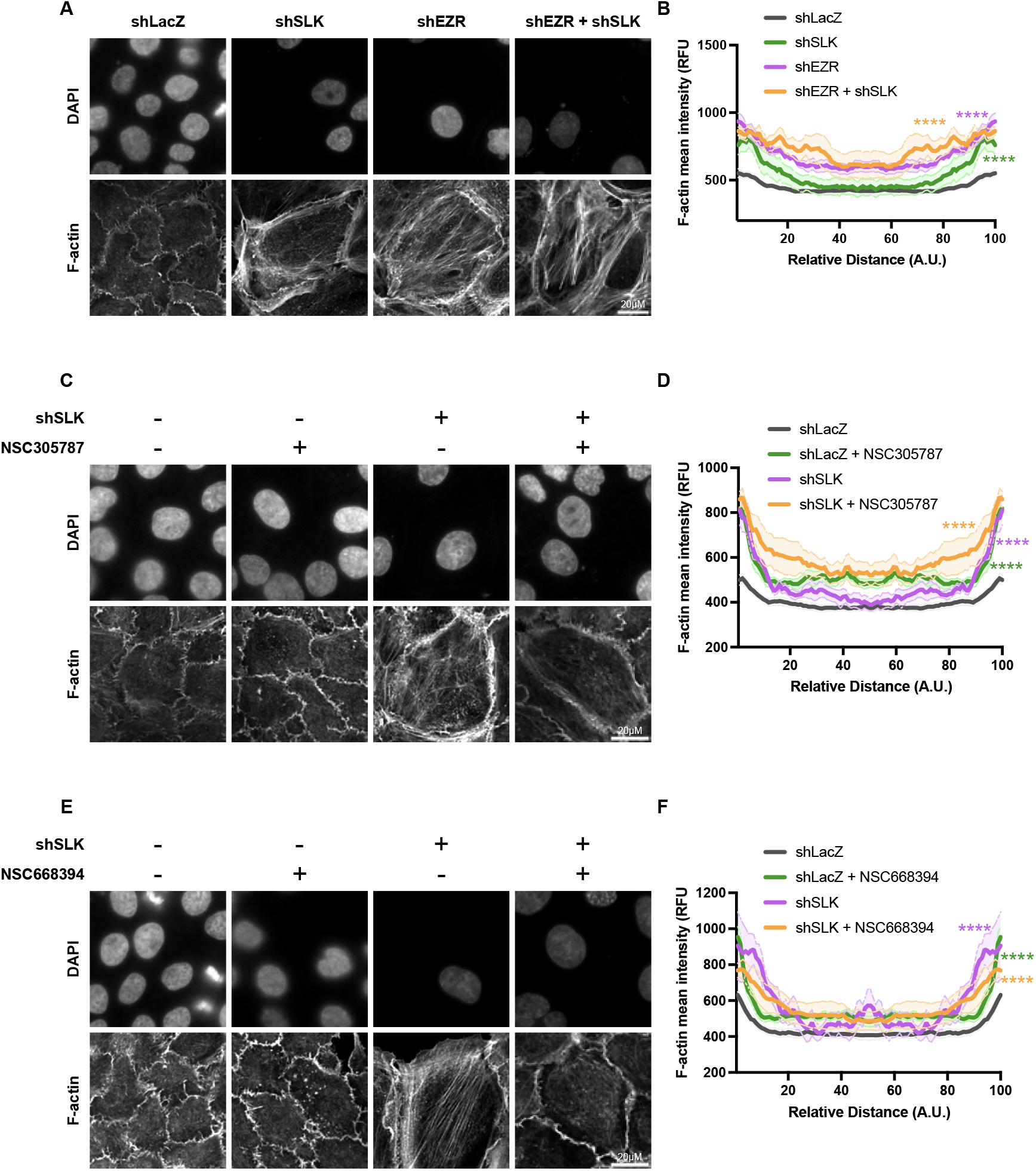
Double-inhibition experiments confirmed kinase-independent facet of SLK-Ezrin cascade in actin regulation. (A)(B) Simultaneous knockdown of SLK and Ezrin increased contractile actin fibers to a similar extent as the knockdown of either SLK or Ezrin alone. (C)(D) Inhibiting Ezrin phosphorylation did not abolish effects of SLK knockdown on contractile actin fibers. Actin signal results were calculated as the average of 12 different lines drawn through each cell at various angles. One-way analysis of variance (ANOVA) and Scheffe’s/Dunn post hoc comparison. Mean ± s.e.m.

Previous research has established that the phosphorylation of ezrin at threonine 567, mediated by the kinases LOK and SLK, is crucial for maintaining microvilli formation and organizing the apical membrane in epithelial cells (*18, 19*). Additionally, the interaction between ezrin and ARHGAP18, a RhoA GTPase-activating protein, modulates RhoA activity in microvilli (*20*). This modulation is vital for maintaining the proper architecture and functionality of microvilli by regulating ezrin phosphorylation and preventing inappropriate myosin assembly. Extending these findings, our results suggest that the SLK-Ezrin cascade may also play a significant role at intercellular junctions, indicating a broader influence on cell architecture. Intriguingly, additional experiments involving the application of ezrin inhibitors in cells with SLK knockdown revealed that the F-actin structure closely mimicked the effects observed with ezrin inhibition alone (Fig 2, C to F). This pattern also suggests that non-phosphorylated ezrin may regulate F-actin in a manner distinct from the mechanisms previously reported, particularly beyond the confines of microvilli structures. These observations contribute to a deeper understanding of ezrin’s role in cytoskeletal organization and highlight potential novel regulatory pathways involving non-phosphorylated ezrin.

### Ezrin mediates contractile actin fibers remodeling through phosphorylation-independent mechanisms

In cellular biology, the protein Ezrin is essential for linking the plasma membrane to the actin cytoskeleton, playing a significant role in cytoskeletal rearrangements. The threonine 567 (Thr567) residue on Ezrin is a critical phosphorylation site, modulating its interaction with F-actin, which is crucial for cell structure and motility (*7, 13, 21, 22*). Phosphorylation of Ezrin at Thr567 by SLK is known to regulate its ability to bind F-actin (*18*). To elucidate the impact of this phosphorylation on Ezrin’s functionality, our study adopted a comparative approach by overexpressing wildtype Ezrin alongside the mutants T567A (non-phosphorylatable) and T567D (phosphomimetic). This strategy aimed to investigate the role of Thr567 phosphorylation in F-actin interaction. Our results revealed that overexpression of wildtype Ezrin, T567A, and T567D consistently reduces the quantity of contractile actin fibers (Fig. 3, A and B). This pattern suggests that the phosphorylation state of threonine 567 influences Ezrin’s ability to stabilize F-actin independently of its activation status. These findings point to an underlying regulatory mechanism that extends beyond the conventional activation-inactivation dynamics of Ezrin. Moreover, simultaneous knockdown of SLK and overexpression of the EZR Thr567 mutants did not result in a significant increase in contractile actin fibers. This observation suggests that effective suppression of contractile actin fiber formation requires cooperative action between SLK and Ezrin (Fig. 3, C and D). It also implies that SLK’s role in modulating F-actin dynamics may involve mechanisms or pathways that do not solely rely on its direct interaction with Ezrin.

**Fig 3.**
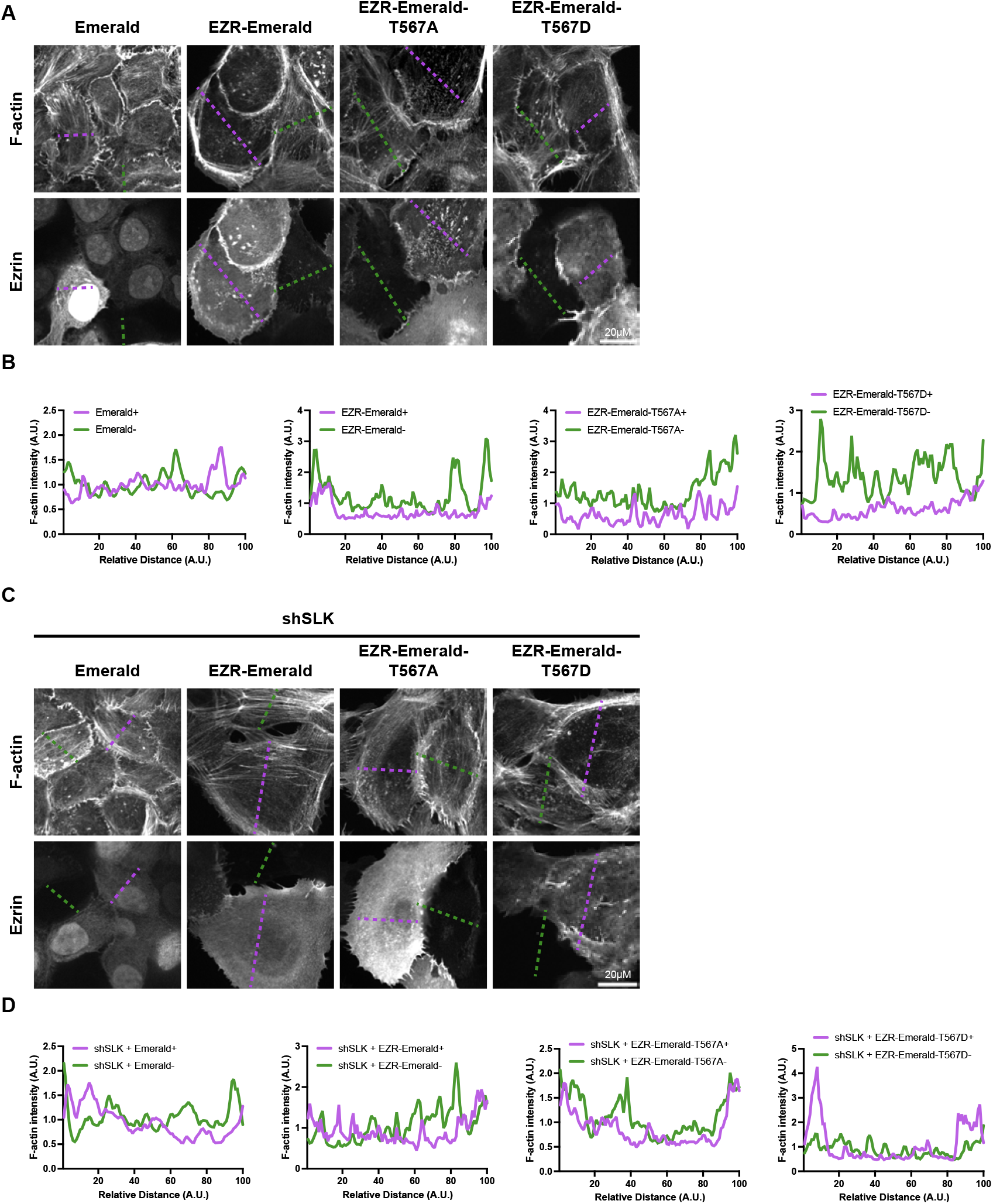
Ezrin reduced contractile actin fibers in phosphorylation-independent manners. (A) Overexpressing various Ezrin mutants in HaCaT cells to investigate their functional impacts. (B) Quantification of F-actin in (A). (C) Simultaneously overexpressing various Ezrin mutants and applying shSLK in HaCaT cells to assess interactive effects on cellular functions. (D) Quantification of F-actin in (C).

### The knockdown of SLK or Ezrin resulted in an increase in contractile actin fibers by altering actin dynamics, rather than by changing actin presentation

Given that the knockdown of SLK or Ezrin leads to an increase in contractile actin fibers, we investigated whether these changes were accompanied by alterations in total actin protein levels. Western blot analysis revealed that these modifications in actin dynamics do not correspond to changes in actin levels (Fig. 4A). This finding is significant, suggesting that the observed effects stem from a reorganization of actin fibers rather than from alterations in actin protein expression. This distinction underscores the regulatory mechanisms that specifically influence actin fiber organization without altering overall protein abundance. Moreover, RhoA, known to be upstream of both SLK and ROCK, plays a pivotal role in the regulatory network affecting the cytoskeleton (*4, 10, 18, 23*). The interplay between these pathways has been extensively documented, often highlighting the close relationship in regulating cytoskeletal rearrangements (*10*). In our previous research, we also observed an antagonistic interaction between SLK and ROCK that influences cell migration processes (*6*). The addition of Y27632 following the knockdown of SLK or Ezrin effectively abolished the previously observed increase in contractile actin fibers (Fig. 4B). This outcome highlights the contractile nature of actin structures mediated by SLK and Ezrin, suggesting that ROCK activity may play a critical role in these effects. Further research is required to elucidate the specific involvement of ROCK signaling pathways in this process (Fig. 4C).

**Fig 4.**
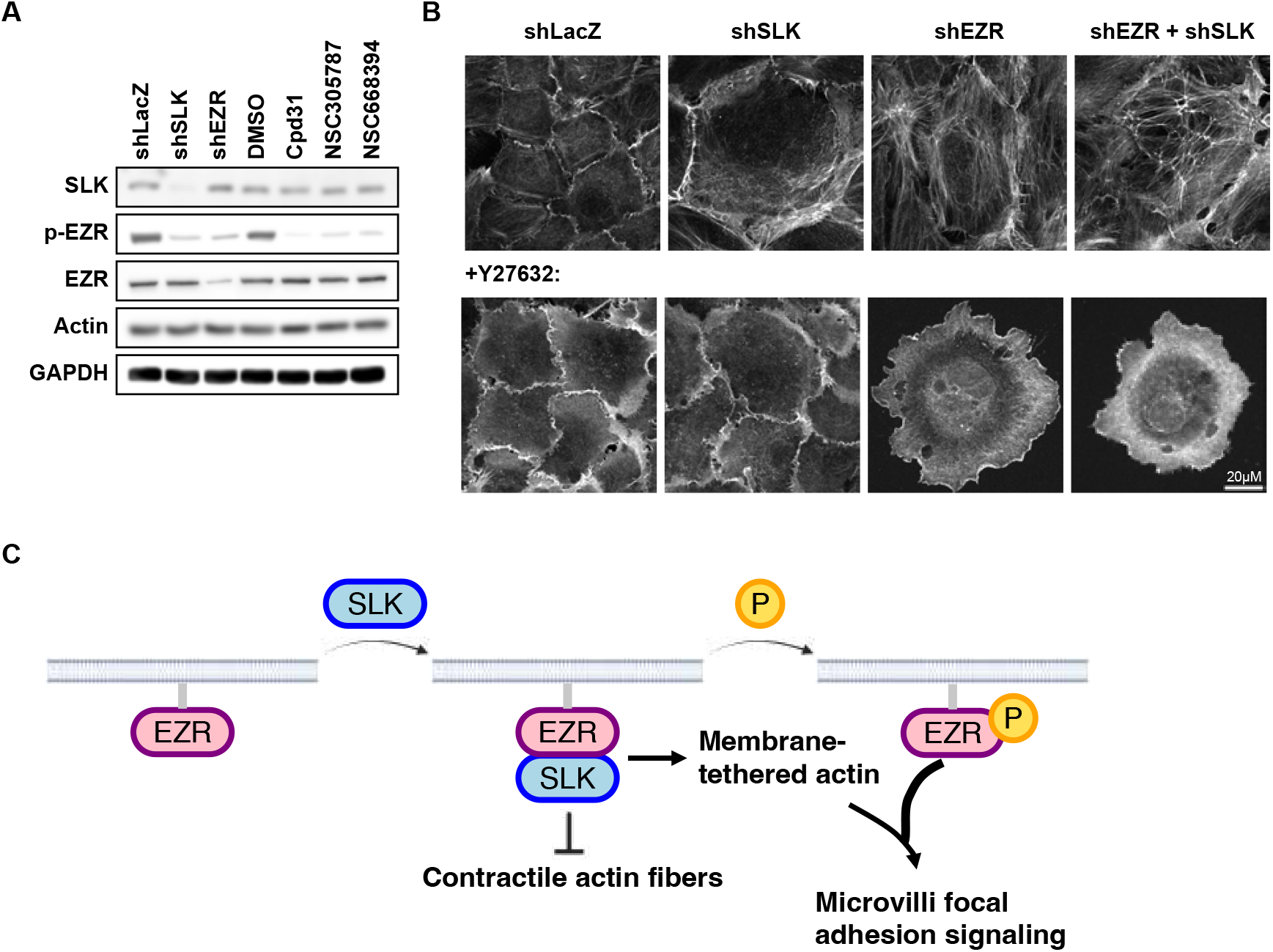
The knockdown of SLK or Ezrin led to an increase in contractile actin fibers by altering actin dynamics, rather than through changes in actin presentation. (A) Western blot analysis is not related to the regulation of actin levels. (B) ROCK inhibition abolished the effects of SLK or Ezrin knockdown on actin fibers, indicating the contractile nature of the actin structures mediated by SLK/Ezrin. (C) The proposed model suggests that unphosphorylated Ezrin can interact with SLK to inhibit the formation of contractile actin fibers and promote the development of membrane-tethered actin. Upon phosphorylation, Ezrin primarily contributes to the formation of cellular structures such as microvilli and focal adhesions.

## Discussion

Our research highlights how SLK and Ezrin orchestrate the cellular framework, particularly the actin cytoskeleton, in ways that extend beyond their known roles in phosphorylation. Our experiments, conducted on different cell lines, demonstrate that the suppression of either protein disrupts normal cytoskeletal structures in a manner not entirely dependent on Ezrin’s phosphorylation state. This suggests that SLK and Ezrin might employ alternative mechanisms, possibly involving structural or scaffolding roles, to influence actin dynamics and maintain cytoskeletal integrity.

Notably, the use of specific inhibitors has allowed us to discern the unique contributions of SLK and Ezrin to the organization of actin networks, particularly in maintaining cell shape and motility. These findings open up new avenues for understanding how cytoskeletal components are regulated, which could be crucial for developing therapeutic strategies targeting cellular motility and adhesion in diseases.

## Materials and Methods

### Cell culture

HaCaT cell lines were provided by the laboratory of J. S. Chia at National Taiwan University, Taipei, Taiwan. Human embryonic kidney (HEK) 293T cells were acquired from the American Type Culture Collection in Manassas, VA, USA. Human umbilical vein endothelial cells (HUVECs) were obtained from Lonza, located in Basel Stücki, Switzerland. All cells were cultured under conditions of 37°C and 5% CO2. HaCaT were cultured in Dulbecco’s modified Eagle’s medium (DMEM; supplied by Gibco, Thermo Fisher Scientific, Waltham, MA, USA and HyClone, Logan, UT, USA). HUVECs were maintained in Endothelial Cell Medium (ECM; Catalog No. 1001, ScienCell). The culture media were enriched with 1% penicillin/streptomycin (P/S) from Gibco and 10% fetal bovine serum (FBS) from BI. Cell passaging was performed using 0.5% trypsin-EDTA from Gibco, and the cells were rinsed with phosphate-buffered saline (PBS) from Corning, New York, USA, within ten passages.

### Generation of knockdown plasmids and overexpression constructs

To silence SLK and EZR genes, HEK293T cells were transfected with shRNA plasmids obtained from the National RNAi Core Facility (NRC; Academia Sinica, Taipei, Taiwan) for the generation of lentiviruses. As a control for knockdown efficiency, shRNA targeting LacZ (shLacZ) was also sourced directly from NRC. Constructs of wild-type Ezrin (pLAS2w.Ezrin-mEmerald), mutant Ezrin T567A (pLAS2w.Ezrin-T567A-mEmerald), and mutant Ezrin T567D (pLAS2w.Ezrin-T567D-mEmerald) were created via In-fusion cloning technology provided by Takara Bio.

### Lentivirus preparation and infection

HEK293T cells were cultured in 6- or 10-cm dishes (Jet Biofil, Guanzhou, China) and transfected with either shRNA plasmids or cloning vectors, pMD2.G and pCMVΔ8.91, using Lipofectamine 3000 (Invitrogen, Thermo Fisher Scientific, Waltham, MA, USA) supplemented with P3000 enhancer (Invitrogen) in Opti-MEM (Gibco). Six hours post-transfection, cells were refreshed with DMEM containing 1% bovine serum albumin (BSA). The lentiviral supernatant was harvested 48 hours post-transfection, followed by centrifugation and concentration using Lenti-X (ClonTech, Takara, Shiga, Kyoto, Japan). HaCaT and HUVEC cells were then infected with the produced lentiviruses, incorporating polybrene (8 μg/ml; Santa Cruz Biotechnology, Santa Cruz, CA, USA) to improve infection efficiency. Selection of transduced cells was carried out using puromycin (2 μg/ml; Sigma-Aldrich, St. Louis, MI, USA) for a duration of 24 to 48 hours.

### Immunofluorescent staining

HaCaT and HUVEC cells were seeded onto 96-well plates (PerkinElmer; #6055300) coated with collagen (100 μg/ml; collagen I, bovine, Gibco). These cells were then transduced with shRNA viruses or viruses carrying overexpression constructs, followed by the selection process using specified antibiotics. Prior to fixation, cells underwent two PBS washes, then were fixed using 4% paraformaldehyde (Sigma-Aldrich) in PBS at room temperature for 15 minutes. Subsequent permeabilization was done with 0.1% Triton X-100 (J. T. Baker, Phillipsburg, NJ, USA) for 10 minutes, followed by a 30-minute blocking step using 5% BSA, all at room temperature. Cells were incubated overnight at 4°C with primary antibodies in PBS containing 1% BSA. For staining, cells were treated with secondary antibodies, phalloidin dyes, and 4′,6-diamidino-2-phenylindole (DAPI; 2 μg/ml; Invitrogen) mixed in 1% BSA in PBS. F-actin was specifically stained using iFluor 594 phalloidin (Abcam, ab176757, 1:1000 dilution). Imaging of the stained cells was performed using a Nikon Eclipse Ti microscope.

### Western blotting

Cells were subjected to viral transduction and subsequently treated for 24 hours with or without a range of pharmacological inhibitors, including PD98059, trametinib, Y27632, and blebbistatin (Sigma-Aldrich). After a rinse with ice-cold PBS, the cells were lysed in radioimmunoprecipitation assay (RIPA) buffer (Cell Signaling Technology, Danvers, MA, USA), augmented with phenylmethylsulfonyl fluoride (PMSF; MD Bio Inc., Rockville, MD, USA) and a mix of protease and phosphatase inhibitors (Thermo Fisher Scientific). The resulting lysates were centrifuged at 15,000 rpm for 15 minutes. The clarified lysates were heated to 95°C for 5 minutes and then applied to an SDS-polyacrylamide gel for electrophoresis. Proteins were subsequently transferred from the gel to a polyvinylidene difluoride (PVDF) membrane (Merck Millipore, Burlington, MA, USA, Immobilon-P PVDF Membrane). This membrane was blocked with 3% BSA in 0.05% Tris-buffered saline with Tween-20 (TBST) (BioLink, Lisle, IL, USA) and incubated with primary antibodies at 4°C overnight, followed by secondary antibodies at room temperature for one hour. Detection of the protein bands was accomplished using enhanced chemiluminescent substrates (T-Pro Biotechnology, New Taipei City, Taiwan), and images were captured with a Bio-Rad Gel Doc2000 (Bio-Rad, Hercules, CA, USA). Quantitative analysis of the proteins was conducted using Image Lab software (Bio-Rad).

---

**Fig. S1.**
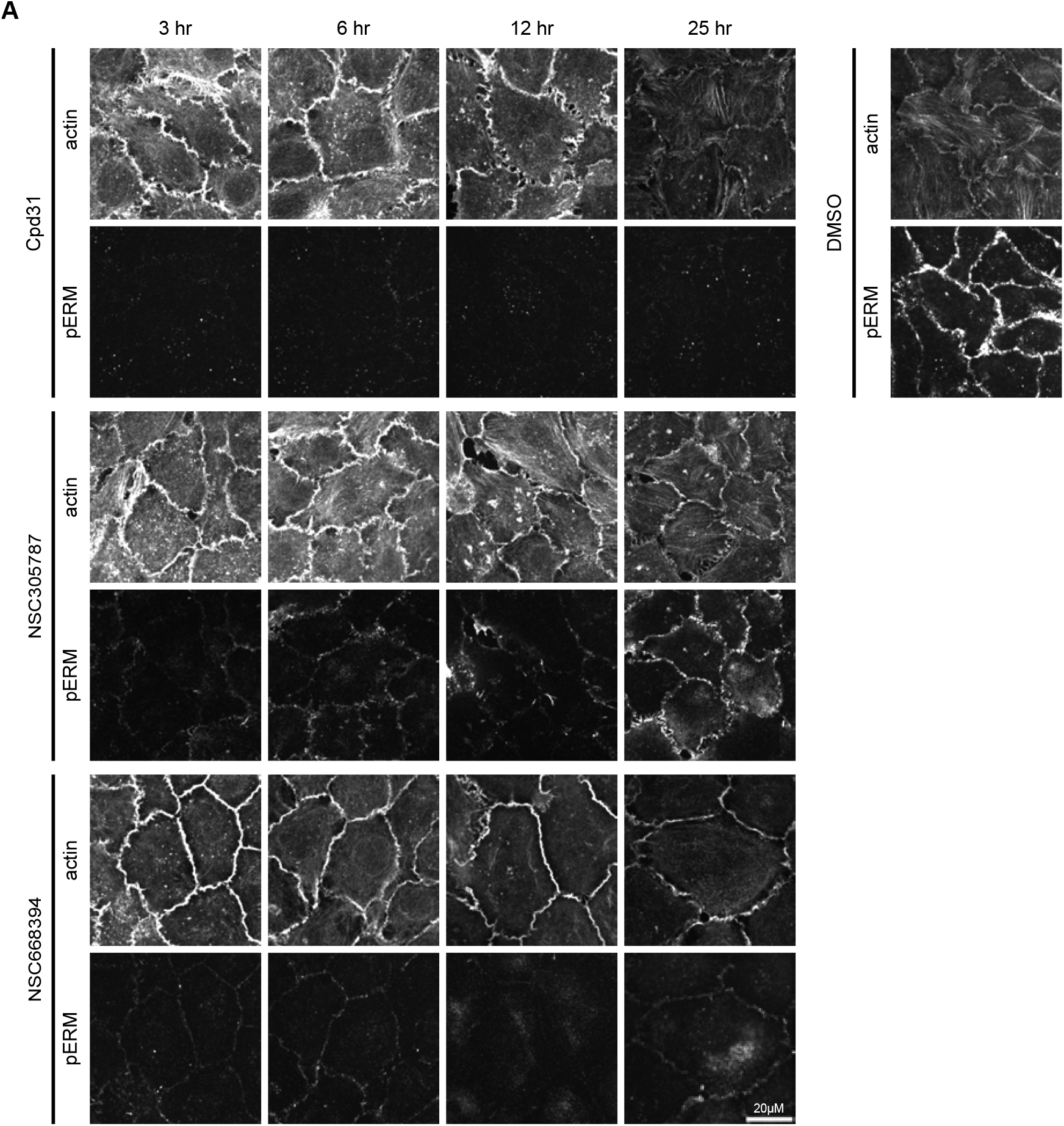
Chronic exposure to these inhibitors at suboptimal concentrations for a duration of 25 hours consistently yielded diminished phosphorylation of ERM (p-ERM), with the observed phenotype of bent cell membranes maintaining stability throughout the period.

## References

1. S. Wagner et al., FAK/src-family dependent activation of the Ste20-like kinase SLK is required for microtubule-dependent focal adhesion turnover and cell migration. PloS one 3, e1868 (2008).

2. S. M. Wagner, L. A. Sabourin, A novel role for the Ste20 kinase SLK in adhesion signaling and cell migration. Cell Adh Migr 3, 182–184 (2009).

3. J. L. Quizi et al., SLK-mediated phosphorylation of paxillin is required for focal adhesion turnover and cell migration. Oncogene 32, 4656–4663 (2013).

4. A. I. Fokin, T. S. Klementeva, E. S. Nadezhdina, A. V. Burakov, SLK/LOSK kinase regulates cell motility independently of microtubule organization and Golgi polarization. Cytoskeleton (Hoboken, N.J.) 73, 83–92 (2016).

5. A. V. Cybulsky, J. Guillemette, J. Papillon, N. T. Abouelazm, Regulation of Ste20-like kinase, SLK, activity: Dimerization and activation segment phosphorylation. PloS one 12, e0177226–e0177226 (2017).

6. L. Y. Yu et al., Synthetic dysmobility screen unveils an integrated STK40-YAP-MAPK system driving cell migration. Sci Adv 7, (2021).

7. A. L. Neisch, R. G. Fehon, Ezrin, Radixin and Moesin: key regulators of membrane-cortex interactions and signaling. Current opinion in cell biology 23, 377–382 (2011).

8. M. Machicoane et al., SLK-dependent activation of ERMs controls LGN-NuMA localization and spindle orientation. The Journal of cell biology 205, 791–799 (2014).

9. L. M. Yin, M. Schnoor, Modulation of membrane-cytoskeleton interactions: ezrin as key player. Trends in cell biology 32, 94–97 (2022).

10. R. Zaman et al., Effector-mediated ERM activation locally inhibits RhoA activity to shape the apical cell domain. The Journal of cell biology 220, (2021).

11. H. Bagci et al., Mapping the proximity interaction network of the Rho-family GTPases reveals signalling pathways and regulatory mechanisms. Nature cell biology 22, 120–134 (2020).

12. R. A. M. Serafim et al., Discovery of a Potent Dual SLK/STK10 Inhibitor Based on a Maleimide Scaffold. J Med Chem 64, 13259–13278 (2021).

13. H. Celik et al., Ezrin Inhibition Up-regulates Stress Response Gene Expression. The Journal of biological chemistry 291, 13257–13270 (2016).

14. A. Mogilner, G. Oster, Cell motility driven by actin polymerization. Biophys J 71, 3030–3045 (1996).

15. R. Ait-Haddou, W. Herzog, Brownian ratchet models of molecular motors. Cell Biochem Biophys 38, 191–214 (2003).

16. R. E. Roberts, M. B. Hallett, Neutrophil Cell Shape Change: Mechanism and Signalling during Cell Spreading and Phagocytosis. International journal of molecular sciences 20, (2019).

17. A. Mogilner, G. Oster, Force generation by actin polymerization II: the elastic ratchet and tethered filaments. Biophys J 84, 1591–1605 (2003).

18. R. Viswanatha, P. Y. Ohouo, M. B. Smolka, A. Bretscher, Local phosphocycling mediated by LOK/SLK restricts ezrin function to the apical aspect of epithelial cells. The Journal of cell biology 199, 969–984 (2012).

19. K. Leguay et al., Interphase microtubule disassembly is a signaling cue that drives cell rounding at mitotic entry. The Journal of cell biology 221, (2022).

20. A. T. Lombardo, C. A. R. Mitchell, R. Zaman, D. J. McDermitt, A. Bretscher, ARHGAP18-ezrin functions as an autoregulatory module for RhoA in the assembly of distinct actin-based structures. Elife 13, (2024).

21. P. Pujuguet, L. Del Maestro, A. Gautreau, D. Louvard, M. Arpin, Ezrin regulates E-cadherin-dependent adherens junction assembly through Rac1 activation. Molecular biology of the cell 14, 2181–2191 (2003).

22. Y. Song et al., Ezrin Mediates Invasion and Metastasis in Tumorigenesis: A Review. Frontiers in cell and developmental biology 8, 588801 (2020).

23. C. Tran Quang, A. Gautreau, M. Arpin, R. Treisman, Ezrin function is required for ROCK-mediated fibroblast transformation by the Net and Dbl oncogenes. EMBO J 19, 4565–4576 (2000).

